# Sonogenetic modulation of cellular activities using an engineered auditory-sensing protein

**DOI:** 10.1101/625533

**Authors:** Yao-Shen Huang, Ching-Hsiang Fan, Ning Hsu, Chun-Yao Wu, Chu-Yuan Chang, Shi-Rong Hong, Ya-Chu Chang, Anthony Yan-Tang Wu, Vanessa Guo, Yueh-Chen Chiang, Wei-Chia Hsu, Nai-Hua Chiu, Linyi Chen, Charles Pin-Kuang Lai, Chih-Kuang Yeh, Yu-Chun Lin

## Abstract

Biomolecules that respond to different external stimuli enable the remote control of genetically modified cells. Chemogenetics and optogenetics, two tools that can control cellular activities via synthetic chemicals or photons, respectively, have been widely used to elucidate underlying physiological processes. These methods are, however, very invasive, have poor penetrability, or low spatiotemporal precision, attributes that hinder their use in therapeutic applications. We report herein a sonogenetic approach that can manipulate target cell activities by focused ultrasound stimulation. This system requires an ultrasound-responsive protein derived from an engineered auditory-sensing protein prestin. Heterogeneous expression of mouse prestin containing two parallel amino acid substitutions, N7T and N308S, that frequently exist in prestins from echolocating species endowed transfected mammalian cells with the ability to sense ultrasound. An ultrasound pulse of low frequency and low pressure efficiently evoked cellular calcium responses after transfecting with prestin(N7T, N308S). Moreover, pulsed ultrasound can also non-invasively stimulate target neurons expressing prestin(N7T, N308S) in deep regions of mice brains. Our study delineates how an engineered auditory-sensing protein can cause mammalian cells to sense ultrasound stimulation. Moreover, owing to the great penetration of low-frequency ultrasound (∼400 mm in depth), our sonogenetic tools will serve as new strategies for non-invasive therapy in deep tissues of large animals like primates.

## Introduction

Approaches that can non-invasively stimulate target cells buried in the deep tissues are highly desirable for basic research and clinical therapy. Currently, different external stimuli including photons, chemicals, radio waves, and magnetic fields have been used to stimulate target cells implanted with stimulus-responsive proteins or nanoparticles^1–4^. However, these strategies suffer from several drawbacks including invasiveness, poor spatiotemporal precision, or low penetration depth, which greatly hinder their potential use in clinical therapy. To overcome these long-standing problems, we aim to use focused ultrasound (FUS) as a stimulus to remotely control cellular activities because it can non-invasively deliver acoustic energy to deep tissues while retaining spatiotemporal coherence^5^.

Ultrasound waves have frequencies greater than those of sound waves that can be heard by humans (>20 kHz). Low-frequency ultrasound waves (<3.5 MHz) are easily transmitted through tissues, including those of bones and brains^6^. Owing to its deep penetrability and spatiotemporal resolution (a few cubic millimetres), ultrasound-based neuromodulation has been tested on cultured neuronal cells and in brains of various model organisms^6–11^. As continuous ultrasound waves or pulsed ultrasound waves of high acoustic pressure are typically needed to activate neurons, neuronal cells are likely to be weakly sensitive to ultrasound stimulus^8,12,13^. To overcome this, gas-filled microbubbles (MBs) that vibrate upon ultrasound excitation have been used as ultrasound amplifiers to enhance their mechanical effects on target cells^14,15^. Recently, lbsen and colleagues used MBs to transduce mechanical stimulation from ultrasound waves to neuronal cells in *Caenorhabditis elegans* and induced behavioural output^16^. The pore-forming cationic mechanotransduction ion channel TRP-4 may be involved in transducing ultrasound stimulation onto MBs attached to *C. elegans*^16^. Although this study verified that ultrasound-mediated neuromodulation is possible, its further development faces major roadblocks, i.e.,, MBs have a short lifespan *in vivo* (<5 min in the blood), and it is difficult to deliver MBs to extravascular tissues^17^. Compared with MBs, gas-filled protein complexes, denoted as gas vesicles, are highly stable both *in vitro* and *in vivo* and efficiently oscillate in response to ultrasound excitation. Different gas vesicle variants can serve as genetically encoded ultrasound contrast reagents to track target microbes or cells by ultrasound imaging^18,19^. However, it is still challenging to express and assemble prokaryotic gas vesicles in mammalian cells^5^. Recently, several groups implanted mechanosensitive ion channels, such as Mscl and Piezo1, into *in vitro* cell culture systems and, with their use, successfully perturbed the cellular membrane potentials of target cells using ultrasound.^20,21^ However, the ultrasound frequencies used in those studies are too high (30 MHz and 43 MHz) to be applicable for *in vivo* use owing to their low penetrability (<5 mm). Therefore, to date, there has been no sonogenetic system that uses low-frequency and low-pressure ultrasound to remotely control activities of mammalian cells that have been genetically modified.

Several mammalian species, including bats and cetaceans, use ultrasound to navigate or communicate. The high-frequency auditory sensitivity and selectivity in echolocating mammals have been attributed to adaptive mechanical amplification in the outer hair cells (OHCs) of their cochlea^22^. Prestin (also known as SCL26A5) is a transmembrane protein residing in OHCs that drives their electromotility and seems to be involved in the ability to hear ultrasound^23–25^. Heterologous expression of prestin endows transfected mammalian cell lines with several of the physiological hallmarks of OHCs, suggesting that prestin may inherently act as an electromechanical transducer^26^. Evolutionary analysis also suggests that prestin is involved in ultrasound sensing of echolocating mammals^23^. The primary sequence of prestin is largely conserved among various mammalian species, although several specific amino acid substitutions that directly affect the electromotility capacity of prestin frequently occur in prestins of sonar mammals but not in those of their non-sonar counterparts^23,24^. Thus prestin probably enhances ultrasound sensitivity in mammals, although how it does so is still unclear.

## Results and discussion

Here we first examined the amino acid sequences of prestin from six non-echolocating species and eight echolocating species. Asn at positions 7 and 308 in prestins of non-echolocating species is frequently replaced with Thr and Ser, respectively, in echolocating species (Fig. 1a). To test whether these apparently evolutionarily driven amino acid substitutions are important to adaptive ultrasound sensing, two mutations N7T and/or N308S were introduced into mouse prestin (hereafter mPrestin). The constructs used for our study were wild-type prestin (mPrestinWT); mPrestin mutants containing a single substitution, mPrestin(N7T) and mPrestin(N308S); and a mutant containing two substitutions, mPrestin(N7T, N308S). Each of these constructs was tagged with the yellow fluorescent protein Venus. Each construct was co-transfected with the calcium biosensor cyan fluorescence protein (CFP)-R-GECO into the human HEK293T cell line. The calcium influx of transfected cells was used as a readout in response to the mechanical stimulation of ultrasound wave. To simultaneously excite FUS and acquire real-time cell images, an ultrasound transducer connected to a waveform generator and an amplifier was placed on top of the live-cell imaging system. This system focuses ultrasound waves to a circle with a diameter of a few millimetres over a monolayer of cultured cells (Extended Data Fig. 1). Using this ultrasound-imaging system, we stimulated cells co-expressing CFP-R-GECO and Venus-mPrestin(N7T, N308S) or co-expressing CFP-RGECO and Venus with a short, low-frequency ultrasound pulse (0.5 MHz, all pulses consisted of 3-sec duration, 2000 cycles, 0.5 MPa unless otherwise noted). Live-cell imaging showed that a short ultrasound pulse of 0.5 MHz was sufficient to evoke calcium influx in cells expressing Venus-mPrestin(N7T, N308S), but not in cells transfected with Venus alone (Fig. 1b; Extended Data Video 1). Quantification of the calcium imaging data indicated that FUS induced a 351 ± 20% (mean ± s.e.m.) increase in the R-GECO fluorescence of Venus-mPrestin(N7T, N308S)—transfected cells (Fig. 1c, right panel). However, FUS only slightly evoked the calcium response in cells that expressed Venus-mPrestinWT (Fig. 1c, middle panel). Cells transfected with Venus alone did not respond to FUS stimulation (Fig. 1c, left panel). These results indicated that heterogeneous expression of Venus-mPrestinWT endowed the transfected cells with a weak ability to sense ultrasound. Substituting Thr at position 7 and Ser at position 308 in the Venus-tagged mPrestinWT substantially improved the ultrasound-evoked calcium response of the transfected HEK293T cells.

**Figure 1.**
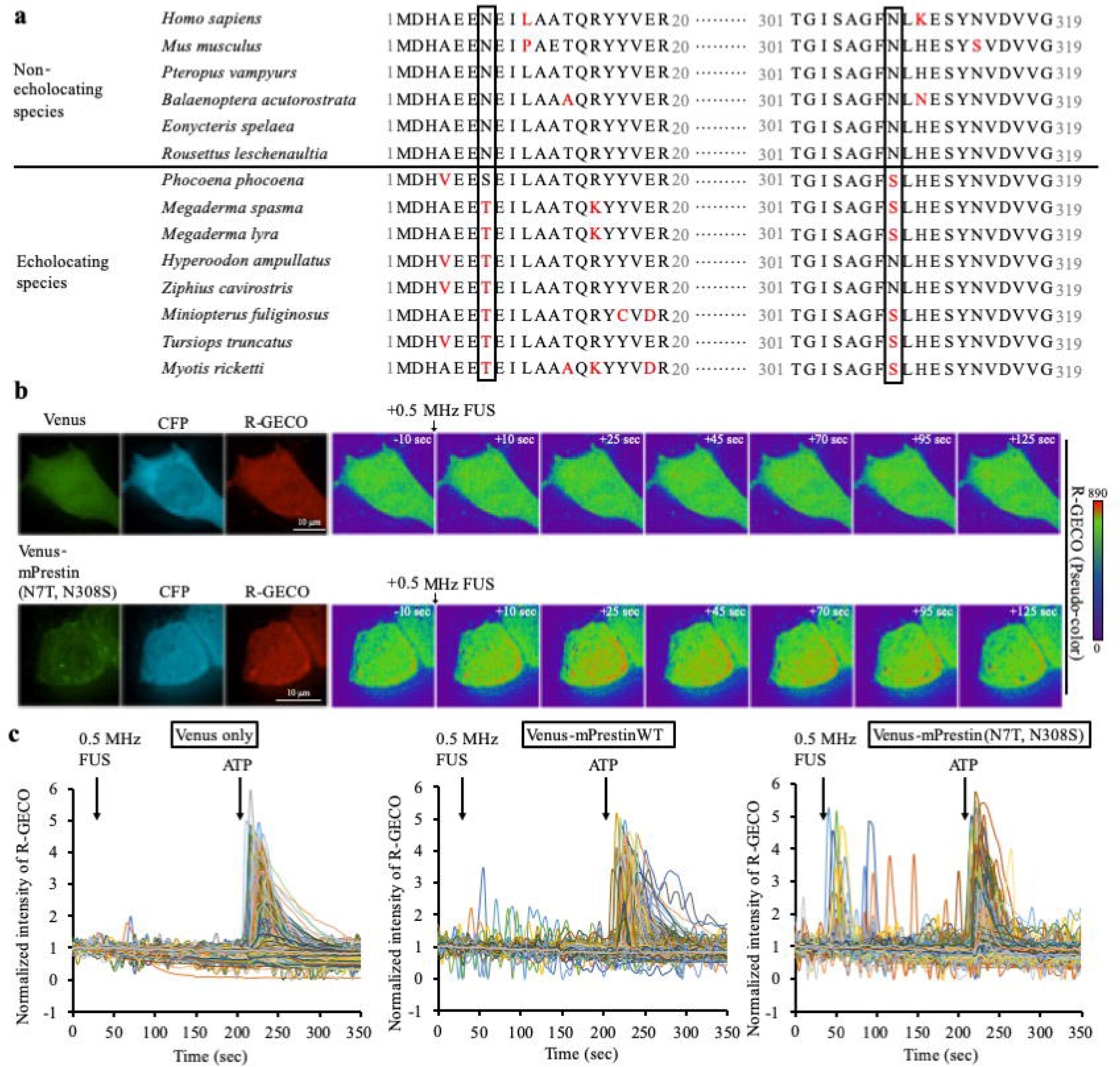
mPrestin carrying the N7T and N308S mutations functions as an ultrasound-responsive protein. **(a)** Sequence alignment of prestins from six non-echolocating and eight echolocating species showing that N7T and N308S substitutions frequently occurred in the echolocating species. Positions 7 and 308 are boxed with the residues located at those positions highlighted in red. **(b)** Excitation of 0.5 MHz FUS evokes calcium responses in cells expressing Venus-mPrestin(N7T, N308S) but not in control cells expressing Venus alone. Cells were co-transfected with the calcium biosensor CFP-R-GECO. The intensity of the R-GECO fluorescence in the cells was monitored by live-cell imaging. Scale bar, 10 µm. **(c)** Time course of R-GECO fluorescence intensity in cells expressing the indicated constructs before and after the 0.5 MHz FUS as described above. ATP treatment (10 µM) served as a positive control to show that the cells could exhibit intracellular calcium flux. Data were collected for 7–36 independent experiments, with *n* = 250 cells per experiment.

To determine the optimal ultrasound frequency/frequencies for cell manipulation, we next comprehensively tested the calcium responses of cells expressing the various Venus-mPrestin constructs to different FUS frequencies between 80 kHz and 3.5 MHz (3-sec duration, 2000 cycles, 0.5 MPa; Fig. 2). Interestingly, cells individually expressing the WT and mutated constructs were sensitive only to 0.5 MHz FUS (Fig. 2). The 80 kHz, 1 MHz, 2 MHz, and 3.5 MHz FUS were insufficient to evoke a calcium influx in the cells (Fig. 2). Upon stimulation by 0.5 MHz FUS, the percentage of ultrasound-responsive cells was 11.29 ± 4.25-fold (mean ± s.e.m.) greater for the Venus-mPrestin(N7T, N308S) group compared with the control group (*p* = 0.024; Fig. 2). Heterogeneous expression of Venus-mPrestinWT, Venus-mPrestin(N7T), and Venus-mPrestin(N308S) only slightly increased the sensitivity of the transfected HEK293T cells to 0.5 MHz FUS (*p* = 0.31, 0.51, and 0.25, respectively; Fig. 2). These results confirmed that 0.5 MHz FUS efficiently evoked a calcium response in cells expressing mPrestin(N7T, N308S) in a frequency-dependent manner.

**Figure 2.**
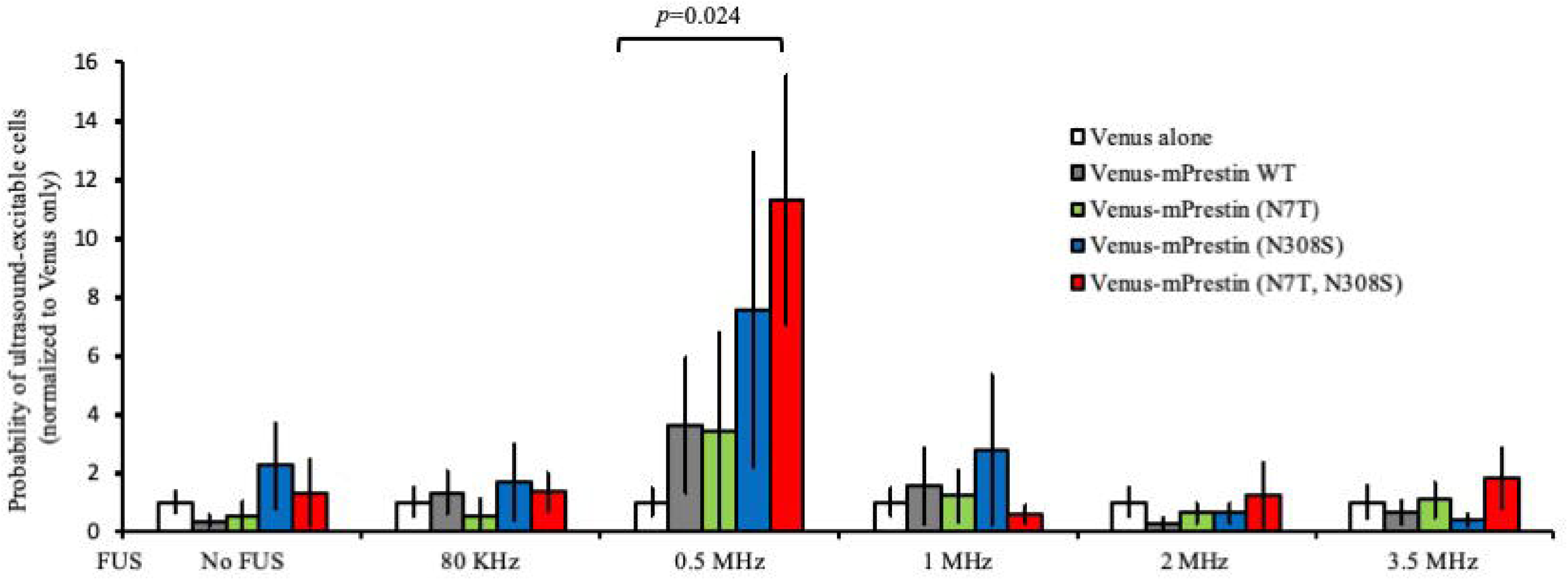
mPrestin(N7T, N308S) enabled an ultrasound-evoked calcium response in a frequency-specific manner. HEK293T cells transfected with one of the indicated DNA constructs were bathed in PBS and stimulated with ultrasound of different frequencies (3-sec duration, 2000 cycles, 0.5 MPa). Data are presented as the relative percentage of cells in each group (expressed as fold-probability) that were excitable by ultrasound after normalisation to that of cells expressing only Venus that were stimulated at the same frequency. The absolute number of cells in each group was 998, 556, 686, 739, 780, 3111, 438, 277, 691, 1484, 1515, 408, 472, 785, 771, 1735, 856, 1571, 1085, 520, 1470, 1050, 1250, 1199, and 605 cells from left to right. Data are shown as the mean ± s.e.m. for 7–36 independent experiments. *P*-values > 0.05 are not shown.

In addition to prestin, Ibsen and colleagues demonstrated that the mechanosensitive ion channel, TRP-4, is required for ultrasound-mediated mechanical stimulation and can modify animal behaviour in the presence of MBs^16^. To test whether TRP-4 can act as an ultrasound-responsive protein, we transfected HEK293T cells with two members of the mammalian TRPC4 family including human TRPC4α (hTRPC4α) use TRPC4β (mTRPC4β)^27,28^. The calcium response of cells expressing hTRPC4α or mTRPC4β tagged with CFP upon FUS stimulation of different frequencies was examined and quantified. The percentage of ultrasound-excitable cells in the in the mTRPC4β-CFP group was 3.29 ± 0.94-fold (mean ± s.e.m.) greater than the control group upon stimulation with 0.5 MHz FUS (*p* = 0.044; Extended Data Fig. 2). Ultrasound of 80 kHz, 1 MHz, 2 MHz, and 3.5 MHz was not sufficient to induce a calcium response in cells expressing mTRPC4β-CFP. Thus, mTRPC4β-CFP is only weakly sensitive to 0.5 MHz FUS. Although its protein sequence is very similar to that of mTRPC4β, hTRPC4α-CFP did not respond to the low-frequency ultrasound stimulation at all (Extended Data Fig. 2). Taken together, the comprehensive examination of ultrasound sensing in cells transfected with different putative ultrasound-responsive proteins shows that mPrestin(N7T, N308S) was the most responsive protein.

We next explored the possible molecular mechanisms that make the two evolutionarily conserved amino acid substitutions important for prestin-dependent ultrasound sensing. Targeting of prestin to the plasma membrane is required for its function^29^. Confocal images of Venus-mPrestinWT and Venus-mPrestin(N7T, N308S) in living cells showed that Venus-mPrestinWT localised to the cytosol and to the plasma membrane, whereas Venus-mPrestin(N7T, N308S) localised exclusively to the plasma membrane (Fig. 3a). Quantification of the relative intensities confirmed that mPrestin(N7T, N308S) exhibited a significantly greater plasma membrane/cytosol intensity ratio than did Venus-mPrestinWT (*p* = 0.003; Fig. 3b). We therefore hypothesised that targeting mPrestin(N7T, N308S) to the plasma membrane is important for its sensitivity to ultrasound. To assess this hypothesis, we introduced a point mutation (Y667Q) that causes prestin to mislocate to the Golgi apparatus into Venus-mPrestin(N7T, N308S). As expected, Venus-mPrestin(N7T, N308S, Y667Q) accumulated at the Golgi apparatus, and its plasma membrane/cytosol intensity ratio decreased significantly (*p* = 1.02E-7; Fig. 3a, b). The mislocalisation of Venus-mPrestin(N7T, N308S, Y667Q) to the Golgi apparatus impaired its ultrasound-sensing ability (*p* = 0.032; Fig. 3c), confirming that plasma-membrane targeting of Venus-mPrestin(N7T, N308S) is required for its response to ultrasound.

**Figure 3.**
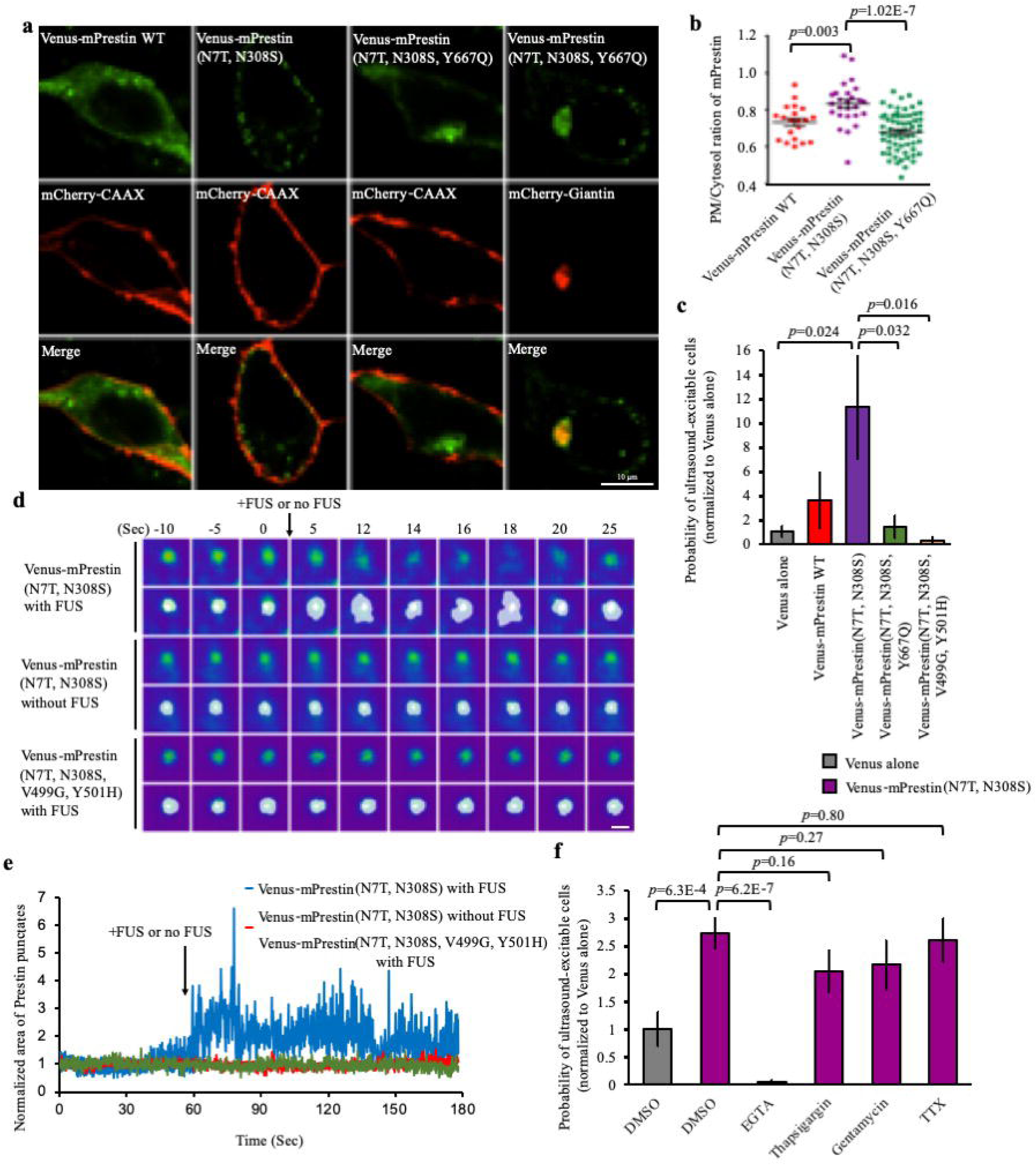
mPrestin(N7T, N308S) puncta oscillate upon FUS stimulation and trigger calcium influx from extracellular pool. **(a)** Representative confocal images of HEK293T cells expressing Venus-mPrestinWT, Venus-mPrestin(N7T, N308S), mCherry-CAAX (a plasma membrane marker), or mCherry-Giantin (a Golgi marker). For each field, the maximum *z*-projection was created from 15 stacks, each separated by 0.3 µm. Scale bar, 10 µm **(b)** Quantification of the plasma membrane/cytosol ratio for the indicated mPrestin constructs. Data are shown as the mean ± s.e.m. for three independent experiments; *n* = 22, 26, and 61 cells from left to right. (**c**) HEK293T cells expressing the indicated constructs were stimulated with 0.5 MHz FUS (3-sec duration, 2000 cycles, 0.5 MPa). Data are presented as in Figure 2. The absolute number of cells in each group was 3111, 438, 1484, 532, and 430 cells from left to right. The data are shown as the mean ± s.e.m. for 6–36 independent experiments. (**d**) Video frames showing the structural dynamics of mPrestin-positive puncta in cells that had or had not been stimulated with 0.5 MHz FUS. The boundaries of the punctate regions are outlined in white. Scale bar, 0.2 µm. **(e)** Area measurements of mPrestin-positive puncta with or without FUS stimulation. **(f)** HEK293T cells expressing Venus-mPrestin(N7T, N308S) were incubated with EGTA (5 mM, 20 min), thapsigargin (100 nM, 30 min), gentamycin (200 µM, 20 min), or TTX (500 nM, 20 min) in DMEM and were stimulated with 0.5 MHz FUS (3-sec duration, 2000 cycles, 0.5 MPa); 0.1% DMSO served as the control. Data are presented as in Figure 1b. The absolute number of cells in each group was 1642, 1238, 1826, 1996, 566, and 772 from left to right. Data are shown as the mean ± s.e.m. for 6–12 independent experiments.

Venus-mPrestin(N7T, N308S) was not evenly distributed in the plasma membrane but was concentrated in punctate regions (Figs. 1b and 3a). HEK293T cells expressing mPrestin(N7T, N308S) exhibit significantly higher number of puncta than cells expressing wild-type mPrestin (p=0.015; Extended Data Fig. 3a). Mislocation of mPrestin(N7T, N308S, Y667Q) at Golgi reduces the number of puncta suggesting puncta formation of mPrestin requires its plasma membrane targeting (*p*=0.032; Extended Data Fig. 3a). Quantification data show the area of mPrestin(N7T, N308S) puncta is 132 ± 6.28 nm^2^ (mean ± s.e.m.; Extended Data Fig. 3b). Prestin self-assembles into oligomers to form bullet-shaped complexes in the plasma membrane^30,31^. To evaluate whether self-association of prestin occurred in these punctate regions, we used fluorescence resonance energy transfer (FRET) to examine the oligomerisation of Venus- and CFP-tagged mPrestin constructs. A greater FRET efficiency was obtained in the punctate regions of cells expressing mPrestin(N7T, N308S) as compared with cells transfected with mPrestinWT (*p* = 0.025; Extended Data Fig. 3c, d), indicating that self-association of mPrestin(N7T, N308S) but not mPrestinWT occurred in the punctate regions. Immunofluorescence staining also showed that mPrestin(N7T, N308S) puncta associated with actin filaments and microtubules (Extended Data Fig. 3e). Next, ultrafast imaging of cells transfected with Venus-mPrestin(N7T, N308S) (imaging interval, 17 ms) was used to observe the real-time behaviour of Venus-mPrestin(N7T, N308S) puncta upon FUS stimulation. Live-cell imaging and quantification showed that Venus-mPrestin(N7T, N308S) puncta oscillated continuously for a few seconds after being exposed to pulsed 0.5 MHz FUS (Fig. 3d, e; Extended Data Video 2). Because several waves of calcium responses were observed after a single FUS pulse in the Venus-mPrestin(N7T, N308S)-transfected cells (Fig. 1c, right), we hypothesised that a short pulse of FUS induced sustained oscillation of Venus-mPrestin(N7T, N308S)-positive puncta that then trigger the calcium response for a few seconds. To address this hypothesis, we found that cellular expression of Venus-mPrestin(N7T, N308S, V499G, Y501H), which prevents the electromotility of prestin without affecting its localisation to the plasma membrane^32^, blocked oscillation of the puncta upon FUS stimulation (Fig. 3d, e; Extended Data Video 2). Moreover, the lack of oscillation found for the Venus-mPrestin(N7T, N308S, V499G, Y501H) puncta significantly attenuated the FUS-mediated calcium response (*p*=0.016; Fig. 3c). Thus FUS-evoked calcium responses are highly dependent on the electromotility and oscillation of prestin puncta in the plasma membrane.

We next determined in which cellular compartment the calcium is stored that is released by Venus-mPrestin(N7T, N308S) upon FUS stimulation. Addition of the calcium chelator ethylene glycol tetraacetic acid (EGTA) in the extracellular space completely inhibited the calcium response in cells expressing Venus-mPrestin(N7T, N308S) upon ultrasound stimulation (*p* = 6.2E-7; Fig. 3f). However, depletion of the intracellular calcium store by thapsigargin did not significantly affect the ultrasound-mediated calcium response (*p* = 0.16; Fig. 3f). Thus mPrestin(NT7, N308S) induced calcium influx from the extracellular space instead of from the intracellular calcium pool after FUS excitation. We speculate that replacement of Asn with Thr and Ser at positions 7 and 308, respectively, in mPrestin enhanced its localisation to the plasma membrane where its oscillations promoted calcium influx from the extracellular pool.

Several mechanosensitive ion channels are activated by high-frequency ultrasound^20,21^. We incubated gentamicin, a pharmaceutical inhibitor of mechanosensitive ion channels, with cells that expressed Venus-mPrestin(N7T, N308S)^33^ and found that this treatment did not significantly affect the calcium response upon ultrasound excitation (*p* = 0.27; Fig. 3f). Thus gentamicin-sensitive ion channels were not involved in the mPrestin(N7T, N308S)-mediated calcium response, which is consistent with results from an ultrasound-inducible system driven by piezoelectric nanoparticles^34^. Ultrasound excites neuronal cells by activating voltage-gated ion channels^6^. To examine the possible involvement of voltage-gated ion channels in our system, cells expressing Venus-mPrestin(N7T, N308S) were incubated with tetrodotoxin (TTX), an inhibitor of voltage-gated ion channels, and then stimulated with 0.5 MHz FUS. However, the percentage of ultrasound-excitable cells transfected with mPrestin(N7T, N308S) was not affected by TTX treatment, indicating that voltage-gated ion channels are not involved in the mPrestin-mediated pathway (*p* = 0.80; Fig. 3f).

To take advantage of the great sensitivity of Venus-mPrestin(N7T, N308S) to ultrasound stimulation, we next developed a sonogenetic system that would allow for stimulating neurons (Fig. 4). Infection of primary cultured cortical neurons with a Venus-mPrestin(N7T, N308S)-containing lentivirus led to the expression of Venus-mPrestin(N7T, N308S) on a neuronal membrane. Moreover, Venus-mPrestin(N7T, N308S) also forms puncta on neuronal membrane (Fig. 4a). For the sonogenetic stimulation of target neurons in deep brain, an adeno-associated virus (AAV) encoding Venus-mPrestin(N7T, N308S) or Venus alone was injected into the VTA brain region. Two weeks later, anesthetized mice were exposed to transcranial pulsed ultrasonic excitation (0.5 MHz FUS, 0.5 MPa, 5 seconds; Fig. 4b). FUS-activated neurons were mapped by imaging the expression of c-Fos (Fig. 4c). Neuronal excitation was triggered by a short pulsed FUS in Venus-mPrestin(N7T, N308S)-transfected mice (*p* = 8.64E-3, Fig. 4c,d). Control mice with Venus alone expression showed no significant c-Fos expression in VTA region (*p* = 0.08, Fig. 4c,d). These results demonstrated that mPrestin(N7T, N308S)-mediated sonogenetics is a flexible and non-invasive approach for sonogenetic control of neuronal activity.

**Figure 4.**
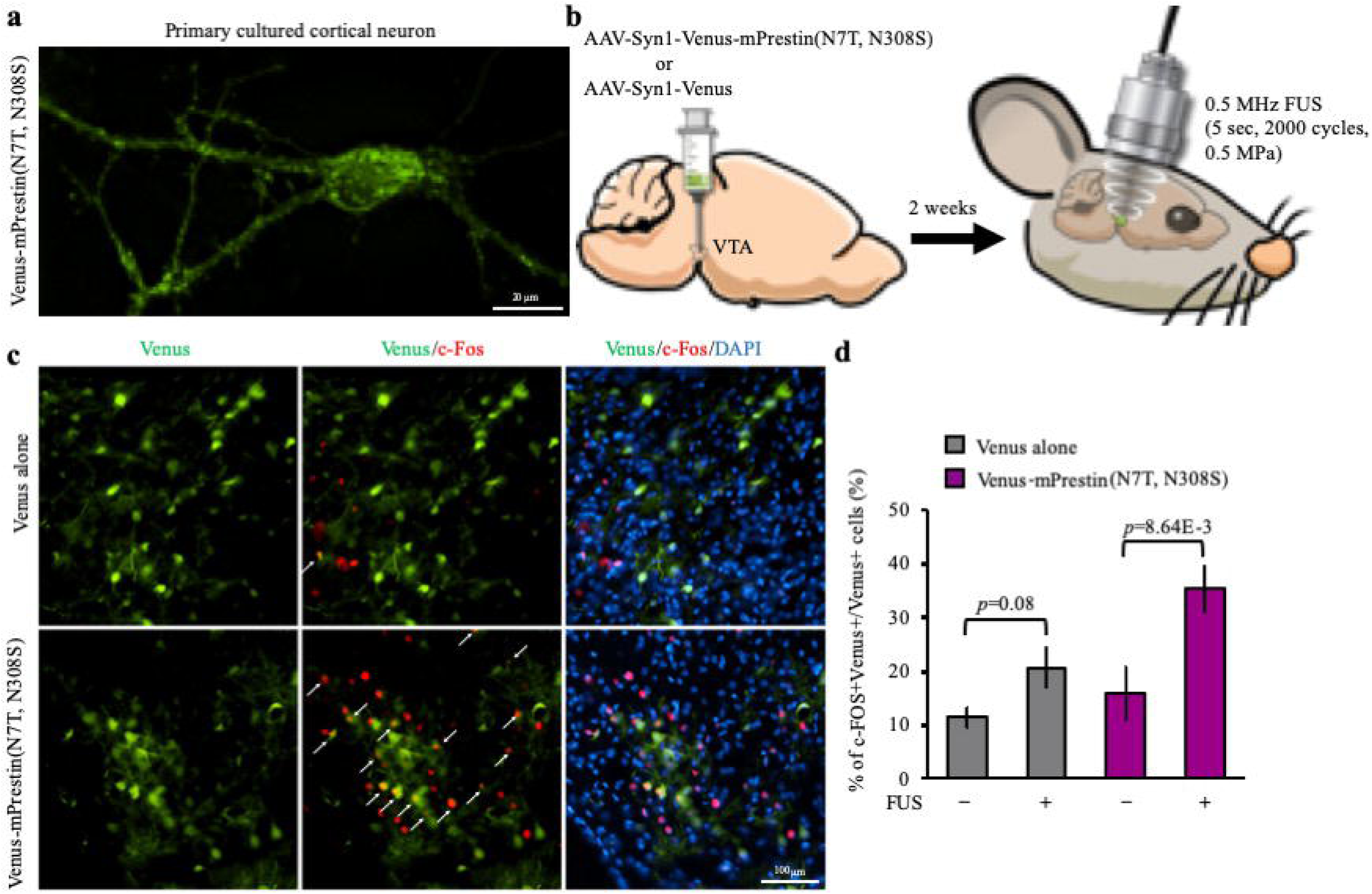
Transcranial FUS stimulation of neuron in mice brain via mPrestin(NT, NS) expression. (a) A representative image of primary cultured cortical neurons expressing Venus-mPrestin(N7T, N308S). The maximum *z*-projection was created from 15 stacks, each separated by 0.3 µm. Scale bar, 20 µm. (b) In vivo experimental scheme for transcranial FUS stimulation of the VTA in anesthetized mice. (c) Representative images of mice brain sections with different conditions. Extensive FUS-driven c-Fos (red) expression was detected in cells expressing Venus-mPrestin(N7T, N308S) after FUS stimulation. Arrows indicate c-Fos+Venus+ cells. Scale bar, 100 µm. (d) Percentage of c-Fos-positive neurons expressing Venus alone or Venus-mPrestin(N7T, N308S) with or without FUS stimulation. Data are shown as the mean ± s.e.m. for 6∼9 different sections from 4 mice per condition.

In summary, we here have introduced two evolutionarily conserved amino acid substitutions N7T and N308S into mouse prestin which enhances its self-association as well as puncta formation in the plasma membrane. These mPrestin(N7T, N308S) puncta highly associate with actin filaments and microtubules in cells. A short pulse of 0.5 MHz FUS induces sustained oscillation of mPrestin(N7T, N308S) puncta with electromotility and evokes several waves of calcium responses in transfected cells (Extended Data Fig. 4). The ultrahigh ultrasound sensitivity of mPrestin(N7T, N308S) allows for non-invasively stimulation of target neurons in deep mice brain by a short pulsed FUS.

Our results raised a fundamental question: why are mPrestin(N7T, N308S)-transfected cells sensitive only to ultrasound of 0.5 MHz? Because we used a short, low-frequency, and low-pressure ultrasound pulse of constant acoustic power (0.5 MPa), it is unlikely that any unexpected thermal and/or mechanical effects were present that would restrict the frequency to 0.5 MHz. We therefore assume that a frequency of 0.5 MHz is simply optimal for stimulation of cells. Cell membranes are able to absorb ultrasound waves and transient cavitation effect occurs in their intramembrane spaces upon ultrasound stimulation^35^. According to simulation and experimental data, 0.25∼0.5 MHz are the optimal frequencies of ultrasound for inducing intramembrane cavitation as well as bio-piezoelectric perturbation^7,36^. Because prestin acts as a piezoelectric amplifier to enhance the electromotile response in OHCs and mammalian cell lines^25,26^, we suggest that the ultrasound-induced intramembrane bio-piezoelectric perturbation may be amplified by mPrestin(N7T, N308S) that then trigger the observed calcium influx. Ultrasound of 80 kHz, which is the peak frequency used by most sonar species^23^, did not efficiently induce a calcium response in Venus-mPrestin(N7T, N308S)-transfected cells (Fig. 2), which suggested that the mechanism(s) of how sonar-responsive species hear ultrasound by auditory organs may not be the same as in our system.

Similar to photon-responsive-proteins and fluorescent proteins, which absorb distinct wavelengths of light and allow for multiplex imaging and optogenetics, mPrestin(N7T, N308S) specifically responds to 0.5 MHz FUS, suggesting that a multiple-frequency system using ultrasound of 1–15 MHz can be developed to non-invasively diagnose regions of abnormal tissues and simultaneously manipulate cellular activities with 0.5 MHz FUS. Moreover, because 0.5 MHz FUS waves cannot be delivered through the air and are rarely used by sonar species, the natural background level for this frequency is expected to be low. Previously developed simulations and experimental data suggest that ultrasound wavelengths of ∼0.60-0.70 MHz would be optimal for transcranial transmission and brain absorption^37,38^, supporting that our sonogenetic system is a promising tool for therapeutic applications involving the brain. Indeed, our *in vivo* results showed that 0.5 MHz FUS efficiently accesses to the deep brain regions like VTA and stimulates target neurons expressing Venus-mPrestin(N7T, N308S) (Fig. 4c,d).

To our knowledge, this mPrestin(N7T, N308S)-based sonogenetic approach is the first system that enables the use of low-frequency ultrasound to efficiently manipulate molecular activities in mammalian cells that are genetically modified. Although heterogeneous expression of mPrestin(N7T, N308S) significantly enhanced the ultrasound sensitivity of HEK293T cells (*p* = 0.0046, 10.18 ± 2.90% for the Venus-mPrestin(N7T, N308S) group; 1.33 ± 0.42% for the Venus-alone group; combined data shown in Figs. 1d and 2f), the percentage of ultrasound-excitable cells in our system needs improvement. A more detailed understanding of how mPrestin(N7T, N308S) sense and amplify ultrasound waves is needed to engineer different prestin variants that are more sensitive to ultrasound. With ongoing development, engineered ultrasound-responsive proteins and sonogenetic systems should become versatile and powerful tools for non-invasively and precisely manipulating activities of genetically modified cells.

## Online Methods

### Cell culture, chemical reagents, DNA constructs, and transfection

Human HEK293T cells were cultured in Dulbecco’s modified Eagle’s medium (DMEM; Gibco) supplemented with 10% (v/v) fetal bovine serum, 5 U/ml penicillin, and 50 µg/ml streptomycin (Gibco). The following Venus- or CFP-tagged mPrestin mutant genes were generated by site-directed mutagenesis: N7T, N308S, Y667Q, V449G, and Y501H. To construct the pLenti-hSyn1-Venus and pLenti-hSyn1-Venus-mPrestin(N7T, N308S), Q5® High-Fidelity DNApolymerase (New England Biolabs) and HiFi® assembly kit (New England Biolabs) were used. The hSyn1-Venus and hSyn1-Venus-mPrestin(N7T, N308S) inserts were PCR amplified from hSyn1-Venus-mPrestin(N7T, N308S) construct. The pLenti-backbone and the insert with a molar ratio of 1:2 (backbone:fragment) were HiFi® assembled to acquire the corresponding lentiviral vectors. For DNA transfection, LT-1 (Mirus) was used according to the manufacturer’s protocol. For inhibitor experiments, gentamicin (200 μM, 20 min, Sigma), TTX (500 nM, 20 min, Abcam), EGTA (5 mM, 20 min, Sigma), and thapsigargin (100 nM, 30min, Sigma) were used. Before ultrasound excitation, HEK 293T cells were incubated with one of the various inhibitors or 0.1% (v/v) DMSO dissolved in DMEM (Gibco) at 37°C. Calcium-free, serum-free medium (Gibco) was used in the EGTA experiment.

### Live-cell imaging

Transfected cells were seeded into a Lab-Tek eight-well chambers (Thermo Scientific) coated with poly-d-lysine (P6407, Sigma-Aldrich) or onto 25-mm cover glasses in six-well culture plates (SPL Life Science) that were similarly coated. Live-cell imaging was conducted using a Nikon T1 inverted fluorescence microscope (Nikon) with a 20×or 60× oil objective (Nikon), a DS-Qi2 CMOS camera (Nikon), and Nikon element AR software (Nikon). The cells were held under a 5% CO_2_ atmosphere at 37°C in an environmental chamber (Live Cell Instrument). The distribution of Venus-mPrestinWT, Venus-mPrestin(N7T, N308S), Venus-mPrestin(N7T, N308S, Y667Q), and Venus-mPrestin(N7T, N308S, V499G, Y501H) in HEK293T cells was observed using a Nikon A1 confocal system with a 100× oil objective (Nikon). Multiple *z*-stack images (0.3 µm between stacks; 15 stacks) were acquired and processed with Huygens deconvolution (Scientific Volume Imaging), and the maximum intensity projections of the images were generated by Nikon element AR software. The plasma membrane/cytosol intensity ratios of the Venus-mPrestin constructs were analysed by Nikon element AR software. Ultrafast imaging was acquired under a Nikon A1 confocal system with a Resonant scanner (Nikon) and 100× objective

### Immunofluorescence staining

HEK293T cells transfected with Venus-mPrestin(N7T, N308S) were seeded on poly-d-lysine-coated Lab-Tek eight-well chambers (Thermo Scientific). Transfected cells were fixed with 4% paraformaldehyde (Electron Microscopy Sciences) at room temperature for 15 min and subsequently permeabilized by 0.1% Triton X-100 (Sigma-Aldrich). After incubation of blocking solution (PBS with 2% bovine serum albumin) for 30 min at room temperature, cells were stained with phalloidin Alexa Fluor 594 (1:100 dilution; Thermo Scientific, A12381) or anti-α-tubulin antibody (1:1000 dilution; Sigma-Aldrich, T6199) for 1 h at room temperature. Goat anti-mouse IgG Alexa Fluor 594 (1:1000 dilution; Thermo Scientific, R37121) were incubated with cells for 1 h at room temperature.

### *In vitro* FUS stimulation

FUS stimulation (acoustic power, 0.5 MPa; 2000 cycles; pulse repetition frequency, 10 Hz; and 3-sec duration) was applied using a single-element FUS transducer (Panametrics). The ultrasound transducer was driven by a function generator (AFG3251, Tektronix) and a radio-frequency power amplifier (80 kHz FUS: 150A100B, AR; 0.5 MHz, 1 MHz, 2 MHz, 3.5 MHz FUS: 325LA, Electronics & Innovation)) to transmit the ultrasound pulses. A water cone filled with degassed water was attached to the ultrasound transducer assembly, after which the surface of the cone was submerged into the culture-dish medium. To record the calcium influx in a cell in real time, the ultrasound transducer was confocally positioned with the objective of the microscope. The transducer was calibrated in the free field in degassed water using a calibrated ultrasound power meter (Model UPM-DP-1AV, Ohmic Instruments Inc.).

### Lentivirus Production

5 h prior to transfection, culture medium of HEK293T cells grown to a confluency of 60% was replaced with 10 ml DMEM supplemented with GlutaMAX (Gibco, Taipei, Taiwan) and 10% FBS (Hyclone, Taipei, Taiwan) containing 25 µM chloroquine diphosphate (Tokyo Chemical Industry, Taipei, Taiwan). HEK293T cells were co-transfected with 1.3 pmol psPAX2 (gift from Didier Trono; Addgene plasmid # 12260), 0.72 pmol pMD2.G (gift from Didier Trono; Addgene plasmid # 12259), and 1.64 pmol transfer plasmids (pLenti-hSyn1-Venus or pLenti-hSyn1-Venus-mPrestin(N7T, N308S)) by PEI (Alfa Aesar; 1 mg/ml polyethylenimine, linear, MW25,000) transfection. The ratio of DNA:PEI was 1:3 diluted in 1 ml of OptiMEM (Gibco). 18 h post-transfection, viral medium was replaced with 15 ml DMEM supplemented with GlutaMAX and 10% FBS. 48 h post-transfection, viral medium was harvested, stored at 4°C and replaced with 15 ml DMEM supplemented with GlutaMAX and 10% FBS. 72 h post-transfection, viral medium was pooled with the 48 h harvest, and centrifuged at 500 × g for 10 min at 4°C. The viral supernatant was filtered through 0.45 µm PES filter (Pall, Taipei, Taiwan), snap frozen with liquid nitrogen, and stored at −80°C.

### Primary neuronal culture and lentivirus transduction

Sprague-Dawley rats were purchased from BioLASCO Taiwan Co., Ltd. Primary cortical neurons were dissociated from dissected cortices of rat embryos (embryonic day 18, E18) and then seeded on poly-L-lysine (Sigma, Saint Louis, MO) -coated bottom-glass dishes (1.2 × 10^6^ cells per dish). On day *in vitro* 0 (DIV0), primary neurons were cultured in Minimum Essential Medium (Invitrogen, Carlsbad, CA) supplemented with 5% fetal bovine serum (Invitrogen), 5% horse serum (Invitrogen), and 0.5 mg/ml penicillin-streptomycin (Invitrogen) under 5% CO_2_ condition. Culture medium was changed to Neurobasal^®^ medium (Gibco, Grand Island, NY) containing 25 μM glutamate (Sigma), 2% B-27™ supplement (Invitrogen), 0.5 mM L-Glutamine (Invitrogen), and 50 units/ml Antibiotic-Antimycotic (AA) (Invitrogen) on DIV1. 10 μM Cytosine-β-D-arabinofuranoside (AraC) (Invitrogen) was added to neurons on DIV2 to inhibit proliferation of glial cells. On DIV3, medium was changed to Neurobasal^®^ medium containing 2% B-27™ supplement, 0.5 mM L-Glutamine and 50 units/ml AA. On DIV6, conditional medium was harvested and half-replaced with fresh Neurobasal/ Glutamine culture medium. Neurons were infected with hSyn1-Venus or hSyn1-Venus-mPrestin(N7T, N308S)-containing lentivirus on DIV7. After overnight incubation at 37°C, the virus-containing medium was replaced with conditional culture medium mixed with equal volume of fresh medium. For further maintenance, the medium was half-changed with fresh Neurobasal/ Glutamine culture medium every 2 days. After lentivirus infection for 7 days, neurons were imaged by a Nikon T1 inverted fluorescence microscope (Nikon).

### Adeno associated virusViral Delivery

The Venus-mPrestin(N7T, N308S) or Venus alone-containing adeno-associated virus (AAV) were packaged by NTU CVT-LS-AAV core. A total of 1 µL of AAV encoding Venus-mPrestin(N7T, N308S) or Venus alone were transcranial injected into the left ventral tegmental area (VTA; bregma: - 3 mm, left: 0.5 mm, depth: 4.2 mm). During the experiment, the animal was anesthetized with 2% isoflurane gas and immobilized on a stereotactic frame. After AAV injection for 2 weeks, the mice were simulated by FUS.

### *In vivo* sonogenetic stimulation of VTA

The AAV transfected mice were randomly divided into four groups: (1) AAV encoding Venus-mPrestin(N7T, N308S) + 0.5 MHz FUS stimulation group; (2) AAV encoding Venus-mPrestin(N7T, N308S) without FUS group(n=3); (3) AAV encoding Venus alone + 0.5 MHz FUS stimulation group (n=3); and (4) AAV encoding Venus alone without ultrasound group(n=3). The 0.5 MHz sonication was applied transcranially at the left brain with the acoustic pressure of 0.5 MPa, 2,000 cycles, and 10 Hz of pulse repetition frequency, sonication duration of 5 sec and one sonication sites. During the experiment, the animal was anesthetized with 2% isoflurane gas and immobilized on a stereotactic frame.

### Immunohistochemistry staining (IHC)

The successful stimulation of mPrestin(N7T, N308S)-transfected cells was verified by c-Fos IHC staining.^39^ The brains of mice were removed were sacrificed at 90 min after 0.5 MHz FUS stimulation. The brains were then sliced into 15-μm sections and incubated into 5% goat serum and PBS for 1 h to block the endogenous proteins. The sections were then incubated in primary rabbit anti-c-Fos antibody (1:1000; SYSY) in antibody diluent for overnight. The sections were then incubated for 1 h in Dylight 594 conjugated anti-rabbit secondary antibody (1:500; GeneTex) in antibody diluent followed by several washes in PBS. The cellular nuclei were labelled by DAPI. Finally, the slides were coverslipped with fluorescent mounting medium and stored flat in the dark at −20°C. The successful transfection of pPrestin was confirmed by the expression of Venus fluorescence protein.

We analyzed the overlap between Venus tagged proteins (Venus alone or Venus-mPrestin(N7T, N308S) and c-Fos by calculating the number of Venus positive cells and Venus/c-Fos double positive cells in different animal groups. Means and s.e.m. were calculated across animals, and all statistics were done across animals.

## Acknowledgments

We thank Dr. Jian Zuo (St. Jude Children’s Research Hospital) for the Venus-mouse PrestinWT construct; Dr. Insuk So (Seoul National University College of Medicine) for hTRPC4α-CFP and mTRPC4β-CFP constructs; and Dr. Takanari Inoue (Johns Hopkins University School of Medicine) for the CFP-R-GECO, R-GECO, mCherry-CAAX, and mCherry-Giantin constructs. Dr. Takananri Inoue (Johns Hopkins University School of Medicine), Dr. Tsung-Han Kuo (National Tsing Hua University), Dr. Hau-Jie Yau (National Taiwan University) for critical reading of the manuscript. This study was supported in part by the Ministry of Science and Technology (MOST), Taiwan (MOST grant numbers 105-2221-E-007-055 and 105-2119-M-182-001 to C.K.Y. 104-2320-B-007-005-MY2, and 106-2320-B-007-004-MY3 to C.P.L. 104-2311-B-007-001, 105-2628-B-007-001-MY3, 107-2628-B-007-001, and 108-2636-B-007-003 to Y.C.L.). Additional funding consisted of a grant from the Program for Translational Innovation of Biopharmaceutical Development-Technology Supporting Platform Axis (grant number 107-0210-01-19-04) to Y.C.L., start-up funding from the National Tsing Hua University to Y.C.L., a grant from Academia Sinica Innovative Materials and Analysis Technology Exploration (i-MATE) Program (grant number AS-iMATE-107-33) to C.P.L., and a grant from National Health Research Institutes (grant number NHRI-EX108-10813NI) to L.C.

## Author Contributions

Y.S.H., C.H.F., and N.H. contributed equally to this work. Y.S.H., C.H.F., C.K.Y., and Yu-Chun Lin designed the experiments. C.H.F. and C.Y.W. programmed the ultrasound system, under the supervision of C.K.Y., Y.C.Chang, S.R.H., Y.C.L., W.C.H., and C.Y.C. performed the cell biology experiments. Y.S.H., C.H.F., N.H., Y.C.Chang, Y.C.Chiang, and W.C.H. quantified the imaging results. Y.S.H., S.R.H., Yen-Cheng Lin, and Yu-Chun Lin generated the DNA constructs. A.Y.W. designed and cloned the lentiviral plasmid, V.G. packaged the lentiviruses, C.P.L. supervised the molecular cloning and lentiviral production processes. C.Y.C. prepared the primary cultured cortical neurons under the supervision of L.C. C.H.F. performed the animal experiments. Y.S.H., C.H.F., C.K.Y., and Yu-Chun Lin wrote the paper.

## Competing interests

The patents of mPresin(N7T, N308S) and relative sonogenetic tools are pending.

## Supplementary information

Supplementary Information is available in the online version of the paper.

## Figure legends

**Extended Data Figure 1.**
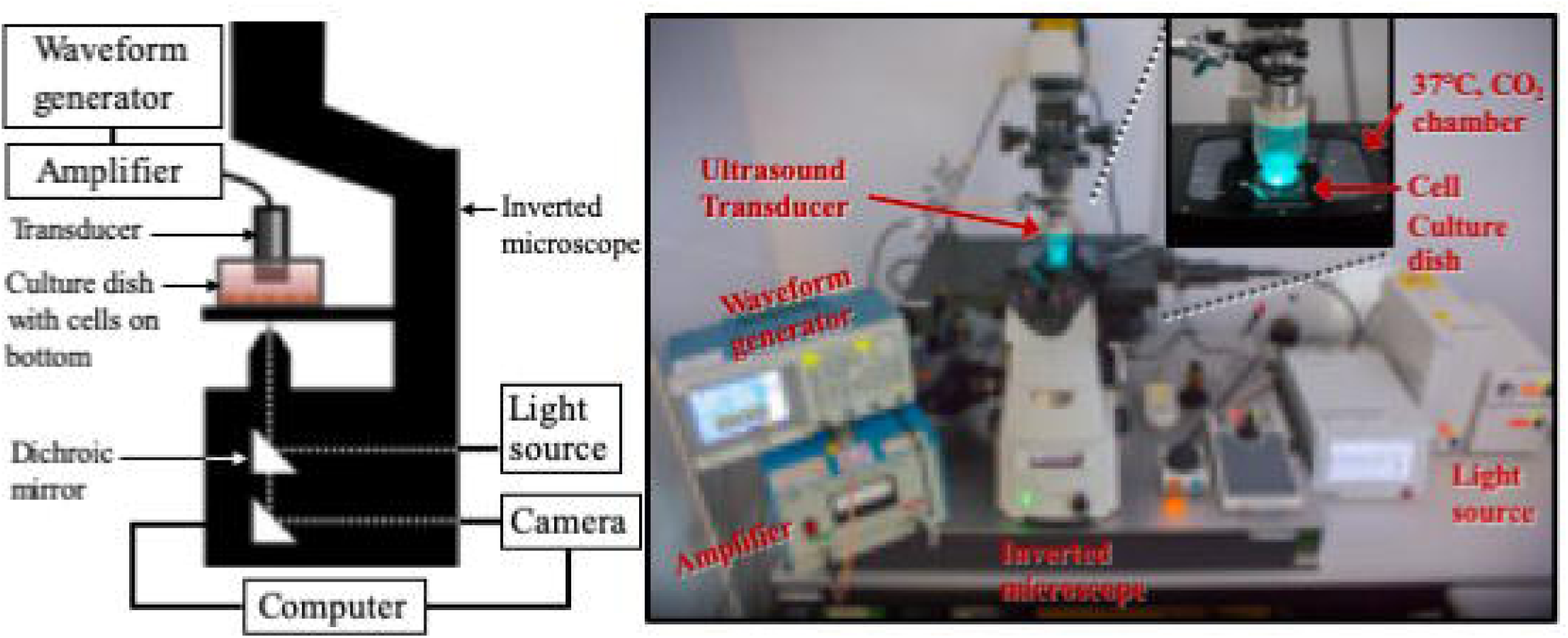
Our computer-controlled live-cell imaging and ultrasound-exposure system. An ultrasound transducer connected to an amplifier and waveform generator was placed in the medium of a culture dish containing a monolayer of cells for FUS excitation. The behaviour of cells upon FUS stimulation in real time was observed through an inverted microscope.

**Extended Data Figure 2.**
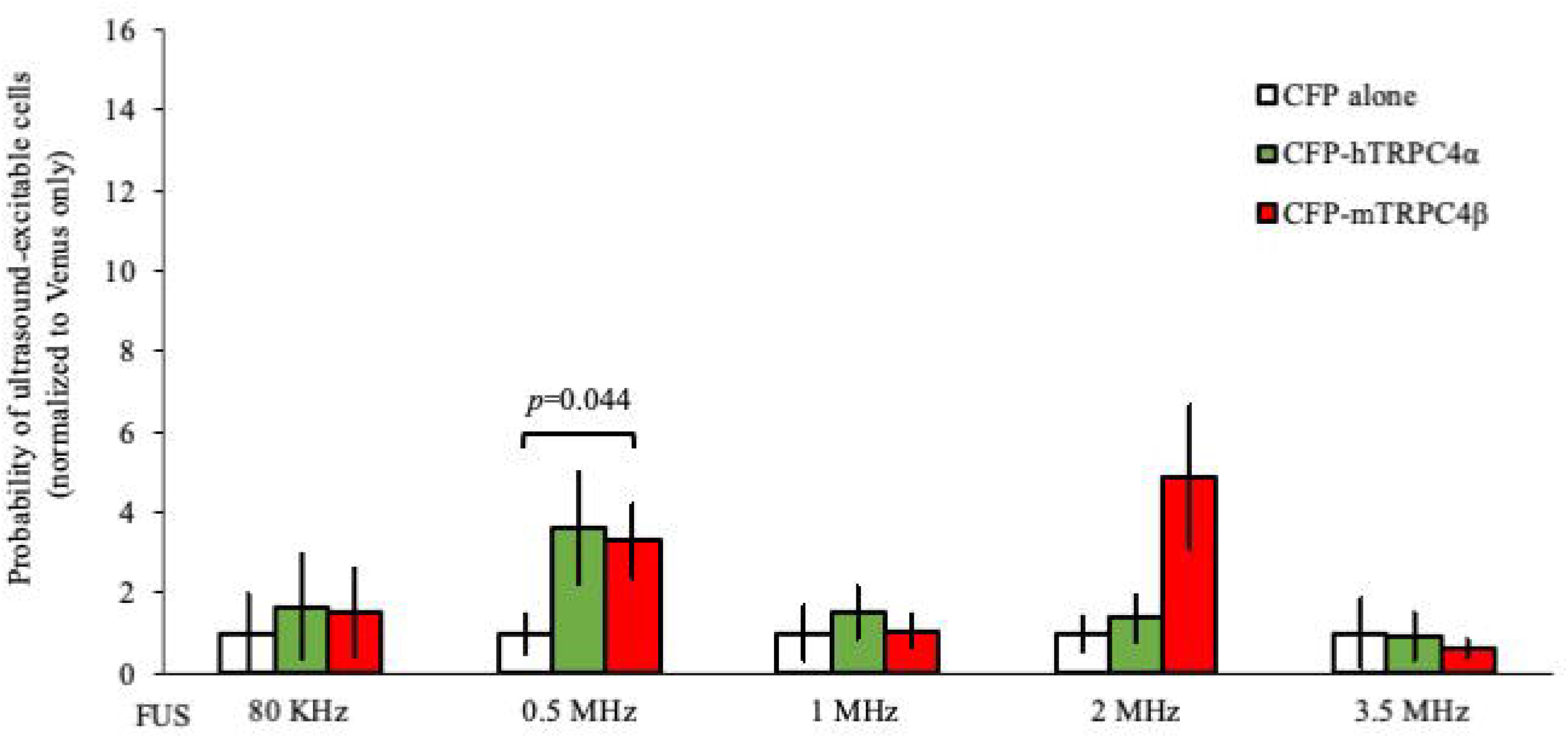
mTRPC4β enabled a weak ultrasound-evoked calcium response in a frequency-specific manner. HEK293 cells transfected with one of the indicated DNA constructs were bathed in PBS and stimulated with ultrasound of different frequencies (3-sec duration, 2000 cycles, 0.5 MPa). Data are presented as the relative number of cells in each group (expressed as fold-probability) that were excitable by ultrasound after normalisation to that of cells expressing only Venus that were stimulated at the same frequency. The absolute number of cells in each group was 1209, 768, 889, 1634, 1665, 1736, 1054, 1035, 1116, 1012, 1000, 960, 1168, 857, and 909 cells from left to right. Data are shown as the mean ± s.e.m. for 7–17 independent experiments. *P* values > 0.05 are not shown.

**Extended Data Figure 3.**
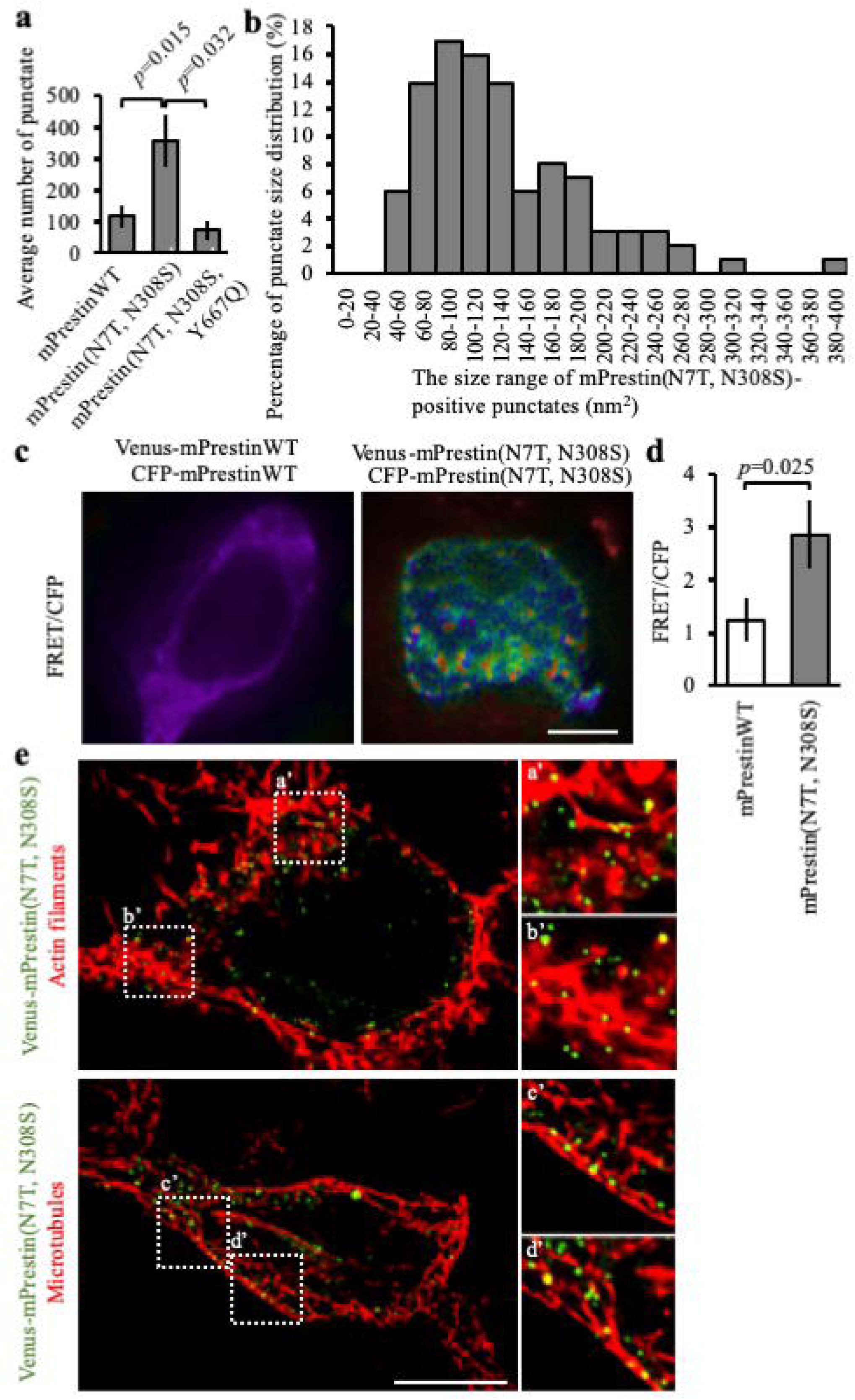
Characterization of mPrestin(N7T, N308S)-positive puncta. **(a)** The average number of mPrestin-positive puncta in cells expressing the indicated constructs. The number of cells counted in each group are 7, 3, and 3 cells from 3 independent experiments. Data are shown as mean ± s.e.m. **(b)** Size distribution of mPrestin(N7T, N308S)-positive puncta. n = 101 puncta from five cells expressing mPrestin(N7T, N308S). **(c)** HEK293 cells transfected with the indicated DNA constructs were imaged by fluorescence resonance energy transfer (FRET). Scale bars, 10 µm. **(d)** Quantification of the FRET/CFP ratios for cells expressing the indicated DNA constructs. The numbers of cells were 25 (mPrestinWT) and 21 (mPrestin(N7T, N308S). Data are shown as the mean ± s.e.m. for two independent experiments. **(e)** HEK293T cells expressing Venus-mPrestin(N7T, N308S) were processed for immunofluorescence with phalloidin (actin filaments) or anti-**α**-antibody (microtubules). For each field, a maximal z projection was created form 15 stacks separated by 0.3 µm. Scale bar= 10 µm.

**Extended Data Figure 4.**
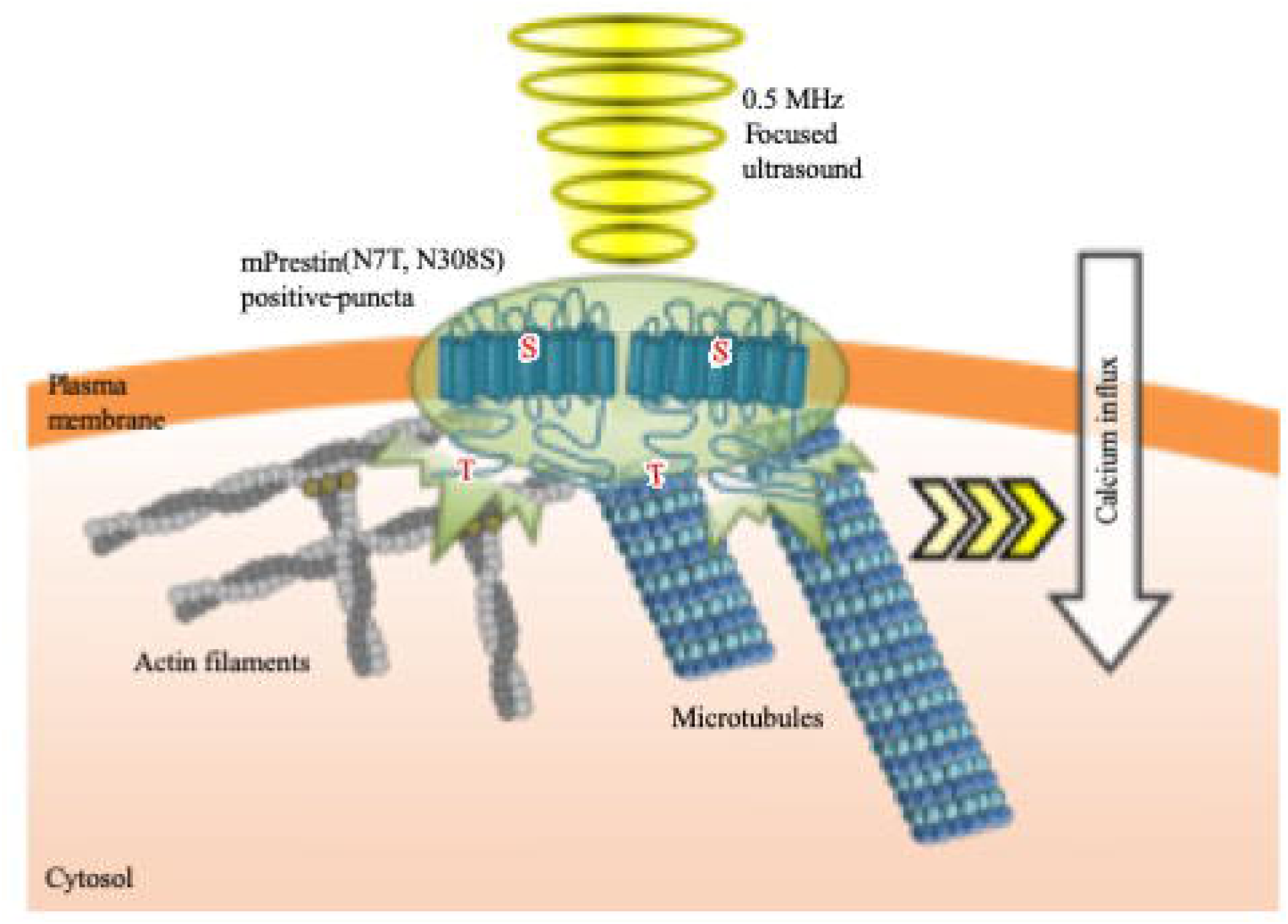
The working model of mPrestin(N7T, N308S)-mediated calcium influx upon ultrasound stimulation. Two evolutionarily conserved mutants N7T and N308S enhance self-assembly of mPrestin in the punctate regions of plasma membrane where they associate with actin filaments and microtubules. The mPrestin(N7T, N308S)-positive puncta with electromotility are oscillated upon 0.5 MHz FUS stimulation (3-sec duration, 2000 cycles, 0.5 MPa) which triggers the influx of calcium from extracellular space.

**Extended Data Video 1 | mPrestin(N7T, N308S) enables ultrasound-evoked calcium response.** Excitation of 0.5 MHz FUS evokes calcium response in cells expressing Venus-mPrestin(N7T, N308S) but not in control cells (Venus alone). The cells co-transfected with a calcium biosensor, CFP-R-GECO, and Venus alone or Venus-mPrestin(N7T, N308S), were excited by 0.5 MHz pulsed FUS (3 sec duration, 2000 cycles, 0.5 MPa). The intensity of R-GECO in cells was monitored by live-cell imaging. Scale bar = 10 µm.

**Extended Data Video 2 | mPrestin(N7T, N308S)-positive puncta oscillated upon FUS stimulation.** HEK293T cells were transfected with Venus-mPrestin(N7T, N308S) or Venus-mPrestin(N7T, N308S, V499G, Y501H). Video showing the structural dynamics of mPrestin-positive puncta in cells that had or had not been stimulated with 0.5 MHz FUS. The boundaries of the punctate regions are outlined in white. Scale bar = 0.2 µm.

